# Lesion Shedding Model: unraveling site-specific contributions to ctDNA

**DOI:** 10.1101/2021.01.28.428297

**Authors:** Kahn Rhrissorrakrai, Filippo Utro, Chaya Levovitz, Laxmi Parida

## Abstract

Sampling circulating tumor DNA (ctDNA) using liquid biopsies offers clinically important benefits for monitoring cancer progression. A single ctDNA sample represents a mixture of shed tumor DNA from all known and unknown lesions within a patient. Although shedding levels have been suggested to hold the key to identifying targetable lesions and uncovering treatment resistance mechanisms, the amount of DNA shed by any one specific lesion is still not well characterized. We designed the Lesion Shedding Model (LSM) to order lesions from the strongest to the poorest shedding for a given patient. By characterizing the lesion-specific ctDNA shedding levels, we can better understand the mechanisms of shedding and more accurately interpret ctDNA assays to improve their clinical impact. We verified the accuracy of the LSM under controlled conditions using a simulation approach as well as testing the model on three cancer patients. The LSM obtained an accurate partial order of the lesions according to their assigned shedding levels in simulations and its accuracy in identifying the top shedding lesion was not impacted by number of lesions. Applying LSM to three cancer patients, we found that indeed there were lesions that consistently shed more than others into the patients’ blood. In two of the patients, the top shedding lesion was one of the only clinically progressing lesions at the time of biopsy suggesting a connection between high ctDNA shedding and clinical progression. The LSM provides a much needed framework with which to understand ctDNA shedding and interpret ctDNA assays.

**Availability:** Binary is available at https://github.com/ComputationalGenomics/LSM

## 1 Introduction

As new cancer sequencing studies continue to become available, it is increasingly evident that tumor evolution and clonal heterogeneity is the rule and not the exception for most cancers [13]. Although a primary tumor can exhibit a set of mutations characterizing the disease, groups of cells, i.e. clones, will acquire new mutations over time creating a disease that looks molecularly diverse at different sites. Once the disease progresses to a metastatic state, it is possible for each lesion to represent a unique subtype of the original disease, harboring their own private mutations [5], that can or can not be sensitive to the administered therapy.

This reality has shifted the paradigm of oncology to greater serial monitoring of the cancer over time with the goal of capturing newly evolved drug targets and possible mechanisms of drug resistance. However, performing serial per cutaneous biopsies on all lesions present in the body at any interval would be overly invasive, and technically infeasible due to the inability to identify all existing lesions at any given time. To confront this challenge, a new era of oncology has begun utilizing liquid biopsies, a noninvasive, low risk method of indirectly sampling tumor components that is amenable to serial sampling of a patient’s entire tumor burden. There primarily exists two types of assays of liquid biopsies that test for different entities: circulating tumor cells (CTCs) and cell-free DNA (cfDNA) [10]. Each have their strengths and weaknesses. Where detecting CTCs can be difficult and costly due to the relatively low number of tumor cells found in the blood, it is a powerful method for early detection, identifying the cancer origin, and generating cell lines on which to test potential therapeutics. However the relative scarcity of CTCs in the blood makes this a more challenging assay. On the other hand, cfDNA assays looking for circulating tumor DNA (ctDNA), which is tumor-derived extracellular DNA found in the plasma, is less costly and provides a means for real time monitoring of patient response and relapse while offering greater potential for discovering new drug resistance mechanisms.

With the growing importance of ctDNA assays, it is increasingly necessary to understand the conditions and factors that cause lesions to shed DNA so as to improve the interpretability and impact of these technologies. The shedding of DNA from the tumor is thought to occur mostly from apoptotic tumor cells, but there is also evidence that they can arise from necrotic cells or be secreted directly into the blood [12]. Interestingly, it has been shown that not all tumors shed ctDNA at the same rate. What influences these different shedding rates remains unknown. There have been several recent studies suggesting shedding rates are influenced by lesion size, i.e. larger tumors will shed more [1]. While other studies suggest that shedding is linked to the molecular profile of the lesion where the more oncogenic a tumor, the higher the rate of shedding [9]. Some studies even suggest that it is the type of cancer that influences the shedding rate, such as higher mortality cancers shedding more than low mortality subtypes. Finally, there are studies that have found that lesions with resistance mechanisms to treatment have the highest shedding rate [7].

Without a clear understanding of the contributory factors to ctDNA levels in the blood, there will continue to be ambiguity as to the significance of detected alterations in cfDNA assays. For instance, if it can be established that larger lesions do in fact shed ctDNA at higher levels, then ctDNA assays need to be adjusted for a potential undersampling of smaller lesions if the read depth is too low. Alternatively if it is found that progressing lesions shed at higher levels, then perhaps alterations with high frequencies in the ctDNA should be prioritized as novel resistance mechanisms as they are more likely shed by those progressing, resistant lesions.

To effectively address the question of what influences tumor shedding rates, there needs to be a means for determining at what levels the lesions are shedding ctDNA. It is for this reason that we developed the Lesion Shedding Model (LSM), a mathematical model of tumor shedding that can determine the relative shedding levels of lesions found in a patient. Here we will describe how the LSM determines the relative shedding levels of lesions and show its relevance in simulated and real patient data.

### 1.1 Lesion Shedding Model

The LSM operates on blood cfDNA and lesion assays to estimate lesions’ ctDNA shedding levels into blood (Figure 1). Patient ctDNA mutation profile represents a mixture of ctDNA shed from all lesions in the body. As there may exist lesions that are not biopsied though are shedding ctDNA into the blood, our framework intrinsically models for missing lesions by subsampling available lesions. This subsampling offers the additional benefits of improving optimization and computational tractability. The LSM takes whole exome or genome sequencing (WES or WGS) data from multiple tissue samples and at least one blood plasma cfDNA sample to develop a mathematical model that identifies the most likely relative contributions of different tumor lesions within each cfDNA sample provided. The LSM operates in two modes: 1) *single time* point mode using the most proximal cfDNA sample to the lesion biopsies, and 2) *longitudinal* mode where each cfDNA sample is analyzed to reveal dynamics in lesion shedding. To develop the LSM, we utilized a combination of simulated cfDNA and lesion data, as well as soon-to-be-released WES data developed as part of the IBM-Broad Drug Resistance Study [3,4,6] that includes patients with multiple synchronous tissue samples taken at autopsy and plasma samples from throughout their treatment course.

**Figure 1.**
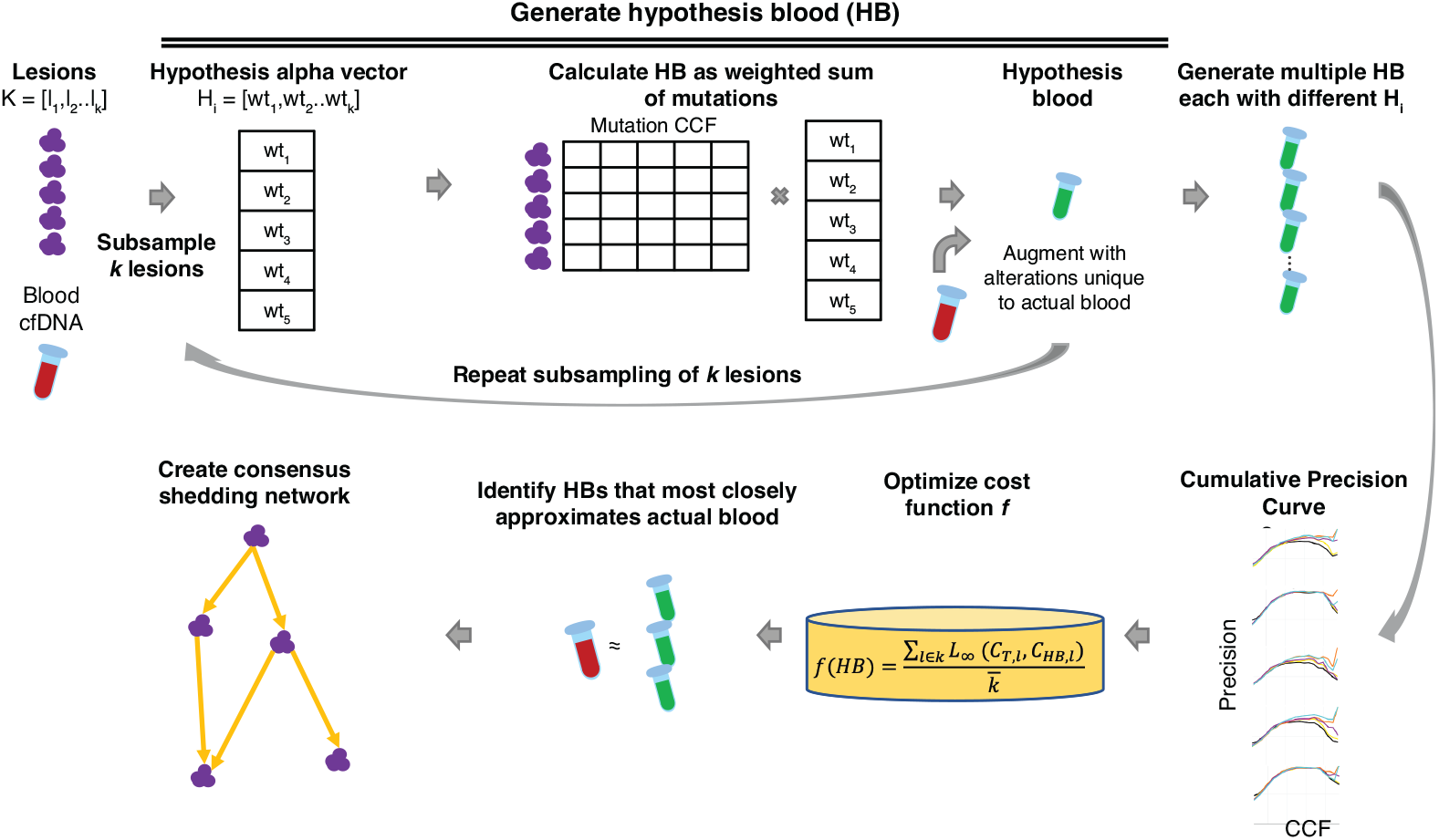
Overview of LSM. Alteration frequencies from blood cfDNA and lesions *K* taken as input. For computational tractability (and robustness), we randomly sub-sample and optimize over up to *k* lesions. This is performed repeatedly to evaluate robustness and satisfies Assumptions 1 and 2. For a given hypothesis blood *HB*, a vector of *k* α, with *alpha* ∈ [0 – 1], *H_i_* is generated. To calculate the *HB*, these *α* are provided as input to a Dirichlet process to generate weights that are multiplied against the respective lesion’s alterations’ CCFs. These weighted CCFs are summed into the *HB*. Multiple HB are generated with different *H_i_*, by default using a grid search to create all possible *H_i_*. Cumulative precision curves of *HB C_HB,l_* and actual blood *C_T,l_* are calculated. Here each plot represents a lesion where the x-axis is the CCF threshold from 0-1 and the y-axis is the precision that the blood sample contains mutations present in the respective lesions. Black line indicates actual blood sample. Colored lines indicate *HB*. We optimized the cost function *f* over all *H_i_* to identify top hypotheses by minimizing the *L*_∞_ distance between *C_HB,l_* and *C_T,l_*. The output is a set of *HB* with lesion-specific weights sorted according to how closely they approximate actual blood. Finally, analysis of all subsamplings yields consensus networks where relative shedding levels are assigned. Lesions are connected by directed edges if the frequency that a source is weighted higher than its target in a significant fraction of subsamplings.

## 2 Methods

**Tumor Shedding Model** We assume the following model: tumors arise from an altered cell, accumulating additional alterations over time. Tumors may give rise to other lesions of similar or divergent compositions, or tumors may continue to arise independently. The model also assumes that tumors may or may not shed ctDNA into the bloodstream over time for a variety of reasons, including apoptosis, necrosis, rapid cell division, etc. Tumors may share similar mutations but their clonal compositions and frequencies can differ between lesions, and thus lesions can be distinguished based on mutational frequency profiles. The blood ctDNA is then composed for all shed ctDNA from all lesions in the body, whether technically assayed or not. The frequency of an alteration in the blood ctDNA can be conceptualized as being a function of the lesion-specific alteration frequency and ctDNA shedding level.

**Terminology** The term *alteration* is applicable to any genetic event including, but not limited to, mutation, single nucleotide variant (SNV), copy number variant, etc. In this manuscript CCF (Cancer Cell Fraction) denotes the fraction of cancer cells bearing an alteration in a cancer sample. For the purposes of our algorithm, CCF and VAF (Variant Allele Frequency) are indistinguishable and the precise method of determining alteration frequencies is outside the scope of this paper. For clarity of exposition we use CCF to represent VAF or CCF and SNV to represent all alterations.

### 2.1 Method Assumptions

**Assumption 1 [Missing Lesion Assumption]** *The model assumes that there may, and likely do, exist lesions that are neither observed nor assayed for biological or technical reasons that shed ctDNA into the bloodstream*.

**Assumption 2 [Non-shedding Lesion Assumption]** *The model assumes that there may be assayed lesions that do not shed, to any significant degree, ctDNA into the blood for a given time period*.

### 2.2 Method Overview

**Input** The LSM receives a MAF file with mutation information, including VAF or CCF. Each patient should include sequencing data from blood cfDNA and lesions *K*. Lesions with < 5% tumor fraction are considered to have poor purity [2] and are excluded from analysis. Sample information is also to be provided in a simple tabular format that minimally provides information on data source (cfDNA or lesion) and sample collection time.

The LSM has four broad phases described below:

1. **Hypothesis Blood Generation** For each patient, generate hypothesis blood *HB* that is a mixture of alterations with an aggregate CCF based on hypothesized lesion shedding levels described by a weight vector *H*.
2. **Target Function** is a family of functions *g_l_*, the goodness-of-fit of *HB* cfDNA profile to lesion *l*, based on cumulative precision curves.
3. **Most Likely Hypothesis** is obtained from multiple runs. For some *k < K*, at each iteration:

a. Randomly select *k* lesions, out of the *K* lesions, for robustness that also naturally addresses Assumptions 1 and 2.
b. For the selected *k* lesions, obtain the optimal hypothesis based on appropriate minimization of the target function (see next section for details) 4. **Consensus Shedding Network** Aggregate top hypothesis from all sub-samplings to calculate a consensus relative shedding level graph.

### 2.3 Hypothesis Blood Generation

The hypothesis blood represents a model of the potential shedding levels of assayed lesions into the blood. By constructing many models with assigned shedding levels, or weights, the LSM can perform a search for the *HB* that most closely resembles the actual blood profile. To satisfy Assumptions 1 and 2 and for computational tractability and robustness, the LSM randomly subsamples and optimizes over up to *k* lesions and this subsampling is performed *S* times. For each subsampling, the LSM constructs a series of hypotheses in the form of assigned shedding levels, *α* ∈ [0,1], to *k* lesions referred to as a hypothesis vector *H*. *α* may be chosen by multiple approaches:

1. lesions assigned *α* randomly by preferred distribution
2. lesions assigned to discrete high, medium, low categories and a random value is drawn from the categories’ specified range. For example:

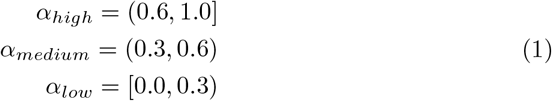
3. lesions assigned *α* from a predefined set, e.g. *α* ∈ [0,1, *increment*], according to a grid search of all possible combinations over *k* lesions. Thus *H* is a vector of *α* of length *k*

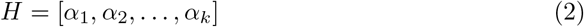

Each *H* is used to create an *HB* sample by providing *H* as input to a Dirichlet process to generate alteration weights that are then multiplied against the respective lesion’s CCFs. For a given alteration *g*, these weighted CCFs are summed across lesions to yield the aggregate *A*(*g*) CCF level in the *HB*.

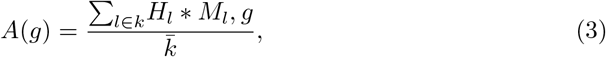

where *M* is a CCF matrix of lesions *l* by alterations *g*, and *M_l,g_* is the set of alterations in *M_l_* whose CCF is ≥ *ct* where *ct* is the CCF threshold below which alterations are filtered out to reduce noise as alterations that are present at low CCF within lesions are likely absent or undetectable in cfDNA. *ct* can be determined by looking at the blood cfDNA CCF level that maximizes *C_T,l_* (see Section 2.4) across most samples in the cohort. Alternatively, *ct* can be adapted per lesion *l* by selecting the CCF that maximizes *C_T,l_* for *l*. In practice, we found *ct* = 0.55 an approximately suitable threshold across many tested samples. The final *HB* is the collection of all alterations’ aggregate CCFs.

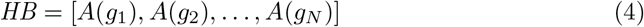

The number of *HB* is determined by the *α* selection strategy.

### 2.4 Target Function Optimization

The LSM identifies the *HB* that most closely approximates the actual blood cfDNA’s representation of alterations found in all assayed lesions. We calculate cumulative precision curves *C* for actual blood cfDNA and all generated HB for each *l* ∈ *k* lesion, *C_T,l_* and *C_HB,l_*, respectively.

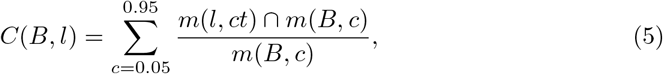

where *B* represents either actual blood or *HB* cfDNA, *c* is the CCF threshold, and *m*(*i,j*) are the number of genes in *i* with CCF ≥ *j*. These precision curves represent how similar the alteration CCFs in the *HB* or actual blood are to the lesions’ CCFs.

The LSM then optimizes a cost function *f* to find the *HB* that minimizes *g_l_*, which calculates the Chebyshev, or *L_∞_*, distance between *C_T,l_* and *C_HB,l_*, over *k* lesions. We selected *L_∞_* distance to minimize the point of maximal divergence between both curves and avoid issues of over-smoothing that may inaccurately identify the best performing *HB*.

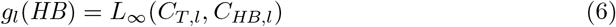

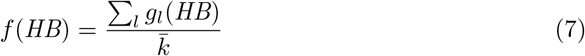

### 2.5 Most Likely HB and Consensus Shedding Networks

For each subsampling *s*, LSM’s output is a set of all *HB* with lesion-specific weights sorted according to *f*. The LSM seeks to minimize *f* and is found to be well-behaved as the distance from the top *HB* solution increases when performing a grid search, i.e. is Lipschitz-continuous. We performed an additional adaptive discrete search of the weight space by using a grid search of the weights centered around the top *HB*’s weights (Appendix Figure 1A,B). Weight ranges were tested ±0.1 along the *H* vector for 50 additional lesion subsamplings. *f* remains well behaved and the lack large shifts in value indicates the solution is stable.

LSM aggregates top *HB* from all subsamplings to calculate a consensus relative shedding level graph. Lesions are connected by directed edges if the source lesion has a higher weight than the target lesion in a high frequency of subsamplings. This frequency can be empirically determined, e.g. conservatively it may consist of two standard deviations above the mean frequency drawn from a null distribution of edge frequencies using randomly selected *HB*s. This produces a directed network indicating each lesion’s relative shedding level. LSM does allow for lesions not to be placed on the graph if an edge cannot be confidently drawn with it. LSM further categorizes lesions into high, intermediate, and low shedding levels using the ratio of outgoing vs incoming edges. High shedding lesions are those where at least 90% of its total edges are outgoing; low shedding lesions are those with at most 10% outgoing edges; and intermediate captures the remainder.

## 3 Results

### 3.1 Simulation Results

We generated simulated cfDNA samples derived from patient data from two gastrointestinal (GI) cancer [8] and one triple-negative metastatic breast cancer [3] patient comparing cfDNA to tissue lesion biopsies to ensure maximal fidelity to actual biological specimens. To first verify the accuracy of the model under controlled conditions, we performed 30 independent simulations where 10 unique cfDNA blood samples were generated from each patient. These unique blood samples were constructed by assigning discrete weights to specific lesions. Three random lesions were given the weights [1.0, 0.6, 0.3] and the remaining lesions were weighted 0.05. Each simulated cfDNA also had 250 random mutations spiked in to add some noise. This simulated blood sample was then analyzed by the LSM. For each patient, *k* lesions were randomly subsampled and optimal hypothesis blood was calculated 200 times. For each subsample, the *H* vector was constructed from weights [0.005, 1.0] at step size 0.25.

We compared the LSM’s consensus directed graph with the assigned lesion weights within each simulation to assess the model’s accuracy. For this graph, the frequency threshold for an edge to be placed between lesions was 1*σ* above the mean frequency 0.45±0.11 taken from a null distribution. We looked at the correlation between each lesion’s fraction outgoing edge and its assigned weight. The expectation was that lesions of higher assigned weight will have a greater percentage of outgoing edges indicating higher predicted shedding level (Figure 2A). The Pearson correlation *r* for simulated patients was *r* = 0.62, *p* < 3 × 10^-44^. The number of lesions for each of the three patients varied from 4-17. We found that while the overall correlation decreased when simulating a patient with higher numbers of lesions, the accuracy in identifying the top shedding lesion remained high and unaffected by the increased lesion count. We also simulated a longitudinal study where at each of five timepoints, three lesions are randomly assigned a shedding weight of 1.0, 0.6, or 0.3 when simulating a cfDNA sample. The LSM is able to correctly order and place those three lesions at each of the timepoints they were assigned (Figure 2B). Though lesions that are shedding at relatively lower levels are more difficult to accurately capture, overall the LSM correctly ordered lesions according to their assigned weight, and it is evident the LSM is most confident in its assignment of the higher shedding level categories.

**Figure 2.**
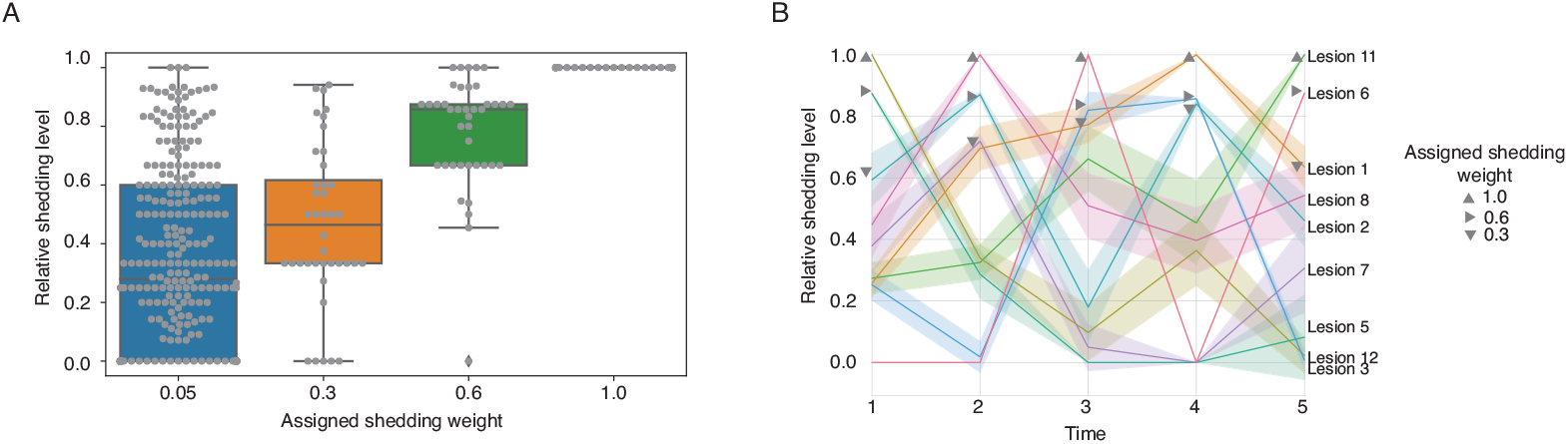
Simulation based controls from 30 independent simulations with 200 sub-samplings derived from three patient samples. A) Boxplot of the mean fraction outgoing edge for lesions grouped by their specifically assigned weights in the simulated sample (x-axis). The Pearson correlation between the assigned shedding weight and outgoing edge fraction is 0.61, *p* < 7 × 10^-33^. B) LSM longitudinal analysis on simulated GI cancer patient. At each timepoint a simulated cfDNA sample was construct with three random lesions assigned a shedding weight of [1.0, 0.6,0.3] and remainder assigned 0.05. Triangles are placed next to the lesions assigned the respective weight. The LSM in-dependently analyzed their relative shedding levels and assessed each lesion’s relative shedding level (solid line) and 95% CI (shaded area).

### 3.2 LSM can capture shedding differences and dynamics in patient samples

From a single cfDNA and multiple lesion biopsy samples, LSM is able to capture the relative shedding differences between lesions. We applied LSM to several patients to understand whether lesions do shed differently and what may be suggested by those differences. We first studied a 53 year old male patient, TPS037, with metastatic BRAF^V600E^ colorectal cancer who was treated with EGFR, BRAF, and PI3KA inhibitors [8]. TPS037 had cfDNA and four tissue lesions biopsies, including two liver, one brain, and one subcutaneous soft-tissue, taken at time of progression. The LSM performed a grid search of the weight space using *α* ∈ [0.005,1, +0.25] discrete increment range over *k* = 4 lesions 200 times. The LSM supports a more detail network view of how lesions relate to each other (Figure 3A) as well as a simplified view to ease interpretation where nodes are discretized into three shedding categories. LSM’s consensus networks reveal the progressing *Liver 1* lesion to be the highest shedding lesion in the blood, suggesting a connection between progression and the cfDNA shedding.

**Figure 3.**
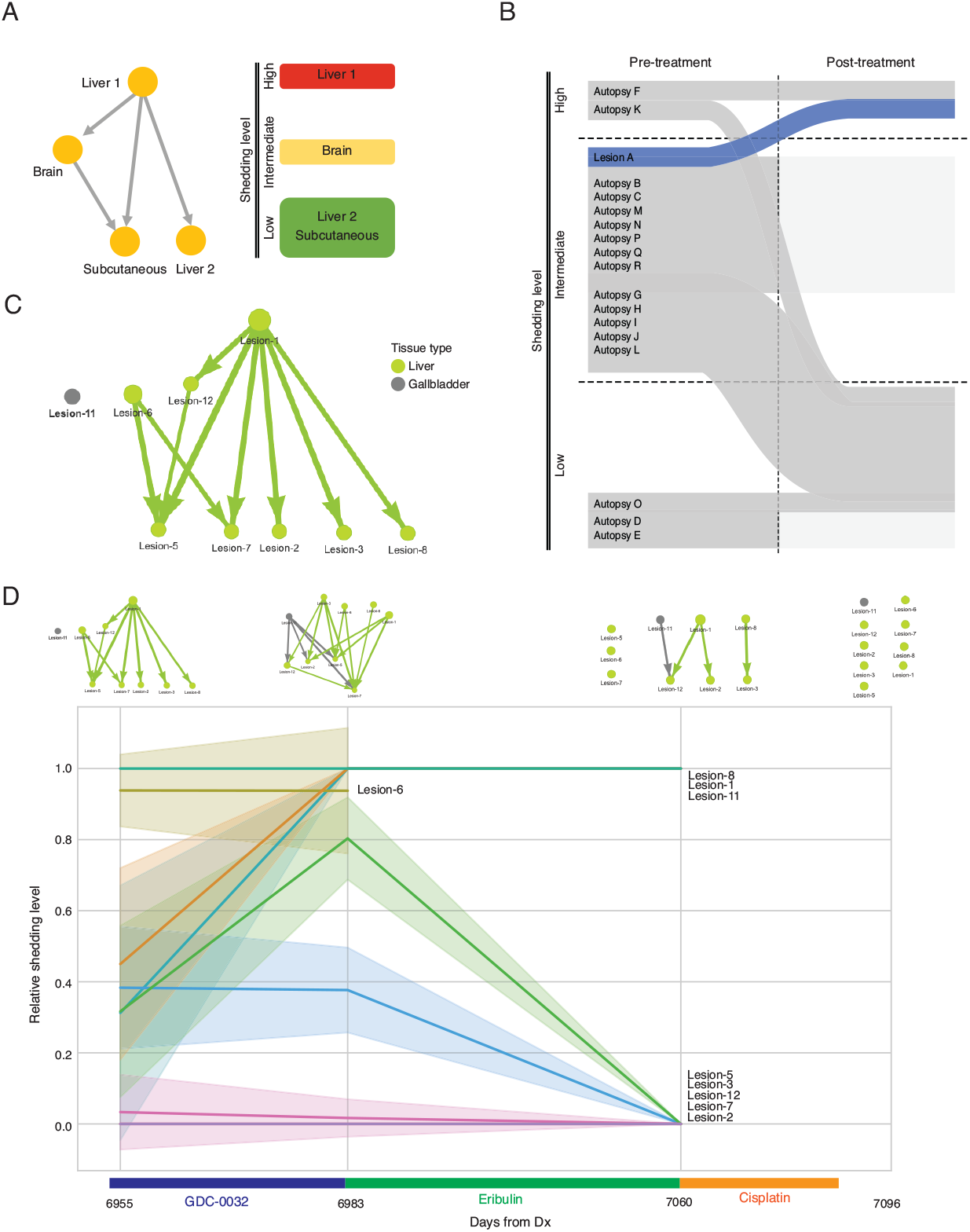
LSM analysis of cancer patients. A) Left - Detailed consensus shedding network for patient TPS037. Right - Simplified consensus shedding figure for patient TPS037. High, intermediate, and low shedding lesions are indicated by the colors red, orange, and green, respectively. B) Subway plot of the longitudinal lesion shedding levels of all lesions in TPS177 pre- and post-treatment with a CDK4 inhibitor. Post-tx lesion A: para-aortic node. Autopsy lesions - B-D: liver, F-H: pancreas, I-J: Mesenteric nodes, K-L: peripancreatic nodes, M-0: Stomach, P-R: Omentum nodes. Blue indicates clinically progressing lesion. Positions where lines terminate indicates lesions that could not be confidently placed w.r.t. other lesions. C) Detailed consensus shedding network for patient MGH-19. Nodes are colored by their tissue of origin. Nodes that are isolated indicates lesions that could not confidently be ordered w.r.t. other lesions. D) Longitudinal shedding plot with each lesion’s relative shedding level (solid line) and 95% CI (shaded area). Above each time is the respective detailed consensus shedding network.

Similarly an analysis of a 58-year old woman with metastatic gastric adenocarcinoma [8], TPS177, revealed a potential connection between tumor progression and high shedding levels. This patient had both pretreatment and postprogression cfDNA biopsies as well as 17 biopsied lesions (one post-treatment and 16 autopsy lesions). With so many assayed lesions, this presented a challenging opportunity to distinguish tumor lesions by their shedding level at each individual timepoint and then connect them to reveal dynamics over the course of treatment. This patient only had lesion-specific progression status information on a single progressing para-aortic lymph node lesion (Lesion A). It was noted in the original study that multiple different FGFR2 mutations were believed to be an underlying resistance mechanism and was identified to be in the liver lesions.

The LSM modeled each timepoint independently, and Figure 3B represents a merging of the simplified shedding representations seen in Figure 3A across timepoints. Similar to TPS037, the LSM found for TPS177 that the confirmed progressing lesion, Lesion A (highlighted in blue), was indeed a top shedding lesion at the time of the postprogression cfDNA sample after it began as an intermediate shedding lesion in the pretreatment sample (Figure 3B). This further suggests that progressing lesions may tend to shed higher into the blood plasma. The LSM also showed that a pancreatic lesion was high shedding across both time points. Lesions from the primary tumor site in the stomach (Autopsy M-O) were all found to be low shedding or indeterminate by the second time point. These stomach lesions were also found in the original study to lack the FGFR2 fusion and had reduced FGFR2 expression and local copy number. All four liver lesions’ shedding levels were found to become indeterminate in the postprogression sample. In this patient, while there does not appear to be a positive correlation between the FGFR2 alteration status and higher levels of shedding in this patient, we once more see the confirmed progressing lesion as a top shedder.

We further tested LSM’s capacity for capturing longitudinal dynamics by applying LSM to a 59 year old female patient, MGH-19, with triple negative metastatic breast cancer [3], who had four cfDNA samples taken over the course of their treatment and nine lesion samples from the liver and gallbladder at time of death. This patient was previously treated with an antibody-drug conjugate, Sacituzumab govitecan (SG), and experienced stable disease according to RECIST 1.1. SG contains a TROP antibody bound to a TOP1 inhibitor. We analyzed the first cfDNA sample, which was the sample closet in time to the end of SG therapy and when the patient began treatment with GDC-0032 (Figure 3C). LSM found Lesion 1, which contained a TOP1 amplification, to be the highest shedding lesion. Lesion 6, which was one of two lesions with a TROP2 copy number amplification and the highest TOP1 amplification amongst all lesions, was also high shedding. The fact that these lesions were amongst the lesions most strongly expressing SG targets may suggest a connection between lesions that should be targeted most efficaciously and high shedders. Figure 3C also highlights that when LSM can not confidently determine a partial ordering for a lesion, it will place that lesion aside as orphaned, as in the case of Lesion 11. We then looked at the remaining timepoints and generated a longitudinal plot of shedding dynamics (Figure 3D). We found that over the course of final three treatments prior to MGH-19’s death there were a great deal of dynamics in the lesion shedding. Lesion 6 along with several other lesions were no longer able to be ordered confidently in the final two months of the patient’s life.

The LSM is able to produce both a detailed consensus directed network as well as a simplified network with lesions categorized into high, intermediate, and low shedding levels.

## 4 Discussion

Liquid biopsies of cfDNA present an important opportunity for real-time, continuous, non-invasive monitoring of patients off and on treatment. These assays offer the means of not only detecting the emergence of new resistance mechanisms but may also provide the basis for understanding the efficacy of a given treatment on specific lesions. As more studies are performed including both tissue and cfDNA, it will be possible to understand if there are shedding biases related to primary disease, cell-of-origin, tissue-of-origin, treatment, etc. that would need to be accounted for when interpreting cfDNA results.

The LSM provides an important first step for understanding the shedding contributions of different lesions into the blood cfDNA. The work presented here is a controlled study on simulated data based on real patient data, as well as providing case studies of two gastrointestinal (GI) and one breast cancer patients. We were able to confirm the LSM’s lesion ordering is robust and not biased by genomic dissimilarity between lesions (see Appendix). In both GI patients, the progressing lesions were found to be top shedding. This finding would suggest giving greater scrutiny to high prevalence alterations in the cfDNA over even known resistance alterations that are found at lower CCF since high prevalence alterations are likely coming from these progressing lesions.

While currently there are relatively few patients with both blood and multi-tissue biopsy data, there exist efforts to address this shortcoming, such as the IBM-Broad Drug Resistance Study [4]. Additionally, extending the LSM to use CTCs as a proxy for lesion biopsies may also address the limited data availability. There are currently assays to extract both CTCs and ctDNA from the same blood sample [11]. Single cell sequencing of CTCs will offer a profile of some subset of lesions that are shedding cells that the LSM can use in lieu of the lesion sample to assess the cfDNA for the relative shedding levels of these “lesions”. The LSM would then be able to monitor ongoing cfDNA samples, which are less expensive and more sensitive than CTC assays, given the relatively few CTCs that are captured in liquid biopsies. Using CTCs as a proxy would provide another mode of data to improve the LSM and the utility of cfDNA assays.

## 5 Conclusion

There are future endeavours to further establish and extend the LSM. These include analyzing additional patients with multiple, synchronous tissue samples and longitudinal cfDNA samples over the course of treatment as part of the IBM-Broad Drug Resistance Study. The increased amount of data will enable us to address the current challenges of ordering lesions that fall into an intermediate category. Currently, the simulated control experiments suggest the top shedding category is the most reliably and confidently assigned. Other shedding level designations should be taken with more caution. Improving the LSM’s ability to resolve intermediately shedding lesions is an area for further study.

The LSM can also be developed to include a likelihood model for the presence of lesions undetected by traditional scans but whose presence explain the blood cfDNA composition. In the same manner and by combining the LSM with models of tumor clonal evolution, we could develop an inference model of the evolutionary trajectory of lesions based on blood cfDNA. With such a model, the LSM could assign new alterations found in the cfDNA as either a likely child of an existing lesion thus showing some kind of evolution or to a new lesion. While there are clearly many important questions still to be addressed, the LSM’s ability to accurately characterize a lesion’s relative shedding level is a vital first step in assigning shedding phenotypes onto lesions from which statistical and machine learning methods can identify the features that explain the mechanisms of shedding and enable cfDNA to be rightly contextualized and its clinical utility dramatically increased.

## 6 Appendix: Model Checking

**Simulation** We developed a module to generate simulated blood cfDNA admixtures to test and verify the accuracy of the LSM under controlled conditions. We simulated a synthetic blood sample per patient where each lesion was given an assigned *α*. A synthetic cfDNA is constructed in similar fashion to Equation (4) using these *α*’s. The simulated blood also has an additional parameter to add random mutations into each sample. This simulated blood sample was then analyzed by the LSM without modification and predicted shedding levels were compared.

**Genetic Bias** To determine whether there is a bias to overweight lesions that are more genetically distinct with respect to others from the same patient, we tested for a correlation the fraction of outgoing edges with samples’ mean genome distance from all other samples (calculated using the Jaccard distance of a binary vector of alterations). The expectation is if there were a bias to assign larger *α*’s to genetically distinct lesions then as the genetic distance increases so to would the fraction of outgoing edges from a lesion indicating a higher assigned *α*. We observed no such relationship in our simulations of patient samples (Appendix Figure 1C) or in real patient data [14]. **Robustness** Lesions are placed in a relative shedding order from high to low by their topological differences, *t*.

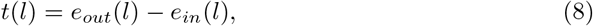

where *e* is the number of edges. To confirm that the relative lesion shedding ordering is robust and stable, we considered the variance of *t* over 50, 100, 150, and 200 lesion subsamplings. The mean of the distribution of the lesion *t* variances over the four sampling amounts is 0.658 ± 1.182 vs a permuted control 13.962 ± 13.746 [14]. The low variance confirms the robustness of the lesion ordering.

## 7 Data and Availability

Patient data was generated and obtained from the following studies: Parikh et al. [8] and Coates et al. [3]. LSM is available at: https://github.com/ComputationalGenomics/LSM.

**Appendix Figure 1.**
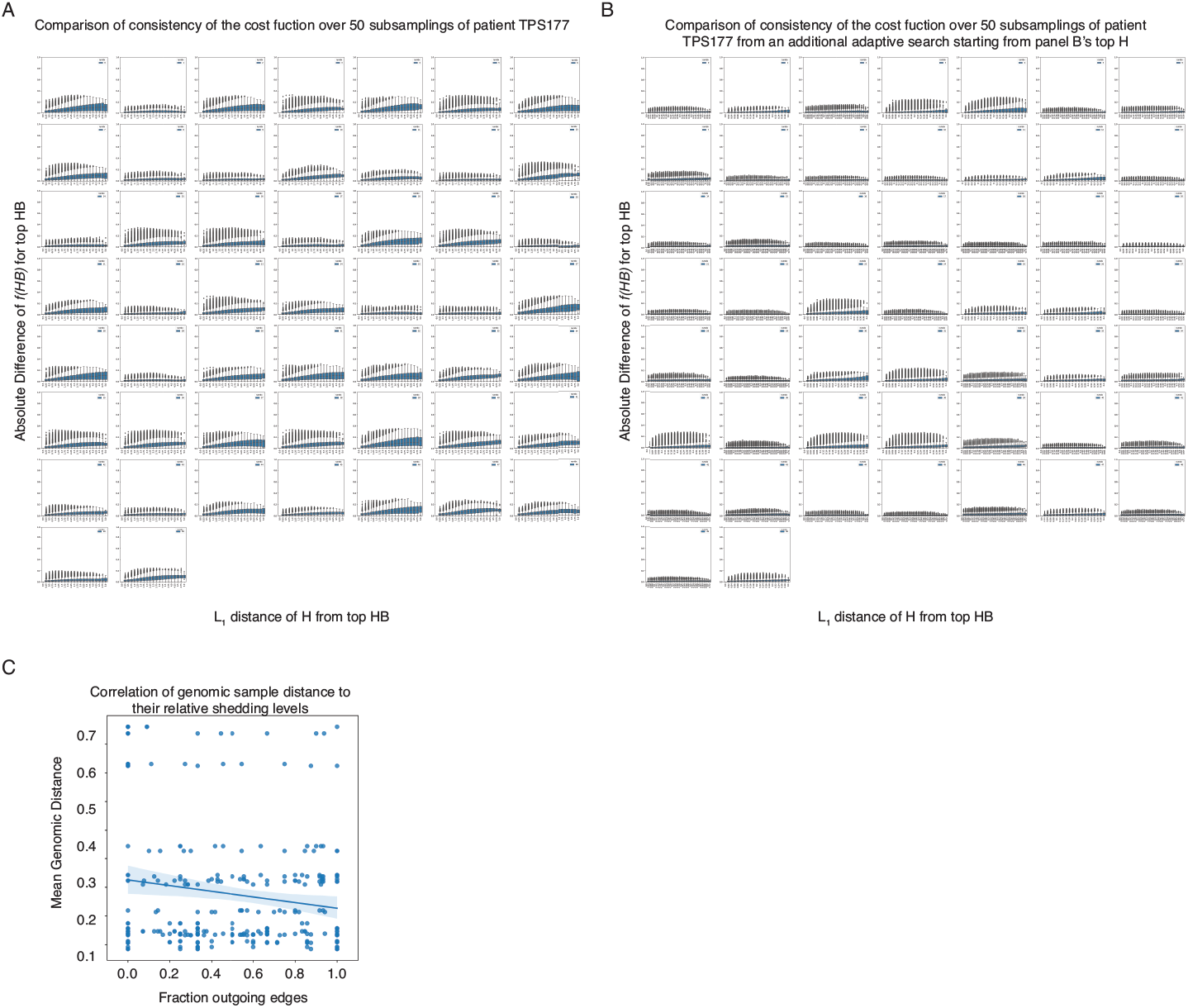
Modeling checking for cost function optimization biases by genomic distances. A-B) Demonstration of consistency of the cost function *f* over 50 subsamplings of GI patient TPS177. A) Each boxplot is a single subsampling with all tested *HBs* exclusive of the top *H*, where the x-axis is the *L*_1_ distance of *H* from the top *HB* and the y-axis is the absolute difference of the *f* (*HB*) from the top *HB*. The cost function f used is well behaved as the distance from the top *f* (*HB*) increases. B) An additional adaptive discrete search of the weight space was performed by using a grid search of the α’s centered on the top *H* from (A). Weight ranges were tested ±0.1 along the top *H* vector for 50 lesion subsamples. *f* remains well behaved and there are no large shifts in values indicating the solution is stable. C) Simulation based controls from 30 independent simulations with 200 subsamplings derived from three patient samples. Scatterplot showing the relationship between the fraction of outgoing edges (x-axis) and the mean genomic distance (y-axis). Each point is a sample in a given simulated run. A linear regression line is drawn.

## 9 Author contributions statement

K.R., F.U., C.L. and L.P. conceived the experiment(s), K.R. and F.U. conducted the experiment(s), K.R., F.U., C.L. and L.P. analysed the results. All authors reviewed the manuscript.

